# A Spatial Whole-Cell Model for Hepatitis B Viral Infection and Drug Interactions

**DOI:** 10.1101/2022.06.01.494377

**Authors:** Zhaleh Ghaemi, Oluwadara Nafiu, Emad Tajkhorshid, Martin Gruebele, Jianming Hu

## Abstract

Despite a vaccine, hepatitis B virus (HBV) remains a world-wide source of infections and deaths, and tackling the infection requires a multimodal approach against the virus. We develop a whole-cell computational platform combining spatial and kinetic models for the infection cycle of a virus host cell (hepatocyte) by HBV. We simulate a near complete viral infection cycle with this whole-cell platform stochastically for 10 minutes of biological time, to predict viral infection, map out virus-host as well as virus-drug interactions. We find that with an established infection, decreasing the copy number of the viral envelope proteins can shift the dominant infection pathways from secreting the capsids from the cell to re-importing the capsids back to the nucleus, resulting in higher viral DNA referred to as covalently closed circular DNA (cccDNA) copy number. This scenario can mimic the consequence of drugs designed to manipulate viral gene expression (such as siRNAs). Viral capsid mutants lead to their destabilization such that they disassemble at nuclear pore complexes, result in an increase in cccDNA copy number. However, excessive destabilization leading to cytoplasmic disassembly does not increase the cccDNA copy number. Finally, our simulations can predict the best drug dosage and timing of its administration to reduce the cccDNA copy number which is the hallmark of infection. Our adaptable computational platform can be utilized to study other viruses, more complex host-virus interactions, and identify the most central viral pathways that can be targeted by drugs or a combination of them.

## Introduction

Whole-cell modeling is a novel approach that aims at simulating the cell cycle and predicting the cell state.^1–3^ These models can be used to either model all essential processes within the cell, or to predict the outcome of specific processes within a realistic cellular environment.^1,4,5^ Whole-cell models can be predictive when subjected to changes in cell environment, mutations, added drugs, and other perturbations that affect multiple network pathways in the cell in a highly inter-dependent manner. They can reveal unexpected side-effects in addition to on-target effects of applied perturbations. Therefore, whole-cell models can provide a platform for next-generation drug development that can replace in-cell and potentially animal studies with cost-effective and more ethical computer simulations, which are not subject to stringent regulatory demands at early stages of the discovery process. Because of the biological relevance of the spatial arrangement of human cells, one of the key ingredients of human whole-cell models is spatial degrees of freedom.^4^

Here we lay the foundation of a computational whole-cell platform for predicting viral infection outcomes in human cells and modeling the effect of different drugs on viral replication in a realistic intracellular environment. We focus on the hepatitis B virus (HBV) that chronically infects 250 million people globally.^6^ Chronic HBV infection, for which there is no cure, can lead to cirrhosis, liver failure, and is the primary cause of hepatocellular carcinoma.^6^ Therefore, there is an unmet medical need to investigate HBV life cycle steps with unknown mechanisms that are related in complex ways through interaction with the host cell cellular machinery.

Like other viruses, HBV exploits the internal machinery of its host cell to replicate. Upon binding to its target, the virus enters the cell in an endosomal vesicle that further releases the capsid into the cytoplasm (Figure 1). The capsid possibly travels along microtubules to reach the nucleus, where its viral genomic content, a relaxed circular DNA (rcDNA), is delivered, possibly by disassembly of the capsid at nuclear pore complexes (NPCs). Inside the nucleus, rcDNA is converted to the covalently closed circular DNA (cccDNA), which is then transcribed to multiple mRNA species, one of which, the viral pregenomic RNA (pgRNA), translates to the core (capsid) protein and the viral reverse transcriptase (RT). PgRNA, RT, and core proteins assemble to form an immature capsid. Conversion of pgRNA to rcDNA within the capsid by RT leads to capsid maturation. Through a mechanism that involves interaction with viral envelope proteins, the mature capsid either re-enters the nucleus for further amplification of cccDNA or is secreted from the cell after acquiring an envelope composed of viral envelope proteins embedded in the host cell lipid membrane.^7^ Additionally, assembly of core proteins alone leads to the formation of empty capsids, that can either be directed to the nucleus or be secreted from the cell.

**Figure 1.**
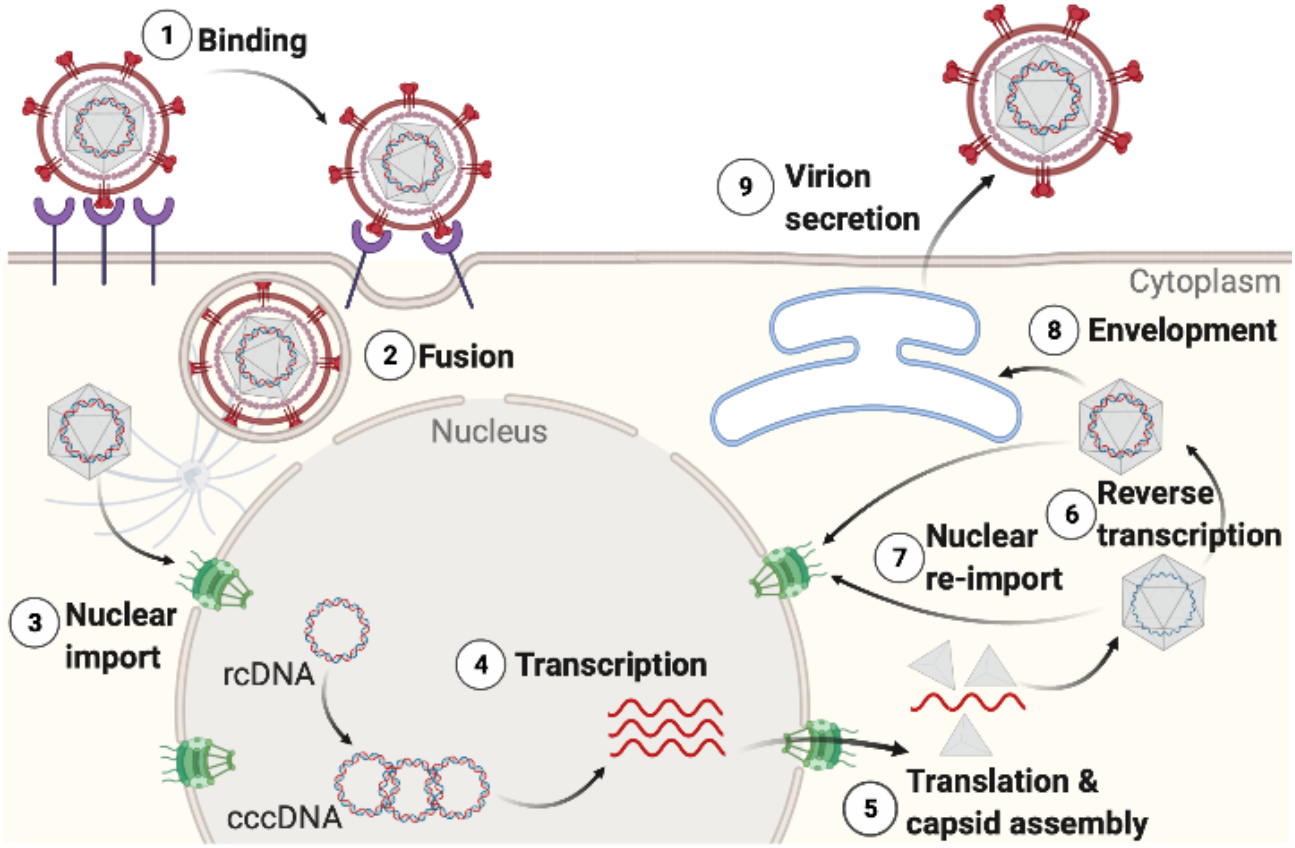
The HBV life cycle. The virion binds to a host cell and the genome-containing capsid (hexagon) is released. The capsid is transported to the nucleus to release rcDNA for repair to convert to cccDNA and transcription. New capsids are assembled and upon maturation, secreted from the host cell (secretion pathway) or re-imported back to the nucleus for further cccDNA amplification (intracellular amplification pathway).

The HBV life cycle has been modeled previously by mathematical and computational approaches. Cangelosi *et al*. developed a multi-scale representation of liver organ by assembling already-published mathematical models for various components such as: the intracellular HBV dynamics and sinusoid-hepatocyte interactions. By introducing gradient variations of key parameters, they studied spatial complexity of hepatic environment on HBV colonisation.^8^ Murray and Goyal^9^ developed a stochastic agent-based model for HBV infection and used it to study various immune response mechanisms for acute infection clearance. Fatehi *et al*.^10^ improved upon the previous model with a more comprehensive reaction network for infection that was solved stochastically. Their results uncovered the presence of different phases in the release of sub-viral particles, and effects of several therapies in different trial phases on HBV infection.

Despite the sophistication of these models, the spatial organization inside the host cell is still missing, namely operating under the assumption that the interior of a hepatocyte is well-mixed. Thus, organelles such as the nucleus and its NPCs, and diffusive or directed transport of capsids within the host are not treated, or lumped into a single reaction rate.^10^ This rate, which depends on the interaction of virus components such as cccDNA, RNA, and core proteins, has been shown to be crucial for the infection outcome, which highlights the importance of spatial resolution in modeling the infection.^10^ Additionally, we have recently shown that other processes such as RNA splicing can be significantly affected by spatial organization within the human cells.^4^

We build a computational whole-cell platform for infection whose response to initial conditions (e.g. capsid counts) and perturbations (e.g. drugs) can be studied to yield a more comprehensive picture of the virus-host cell interaction. We took a data-driven approach to develop a model of a realistic cellular environment, by integrating multiple available experimental data. We then developed a kinetic model that describes the viral infection cycle and complemented it with a diffusion model describing how various species diffuse in different regions of the modeled cell. Both kinetic and diffusion models were parametrized with experimentally determined values. Finally, we performed reaction-diffusion master equation simulations on the entire HBV life cycle within the *in-silico* hepatocyte for 10 minutes of biological time to explore two key questions as our first application of the model: How does the expression of viral envelope proteins on the endoplasmic reticulum (ER) membrane affect the decision process for capsids being released vs. being reimported to the nucleus to further amplify cccDNA? And what is the optimum dosage and timing for HBV capsid inhibitor drugs as predicted by the model?

The development of this platform can help virologists and drug developers interested in HBV infection to better design experiments by testing hypotheses with the model. Such a virtuous cycle of whole-cell model and experiment could lead to a comprehensive understanding of the ‘hotspots’ in viral infection that need to be treated by a multimodal approach.

## Results

### Spatially-resolved model of a hepatocyte

Following our previous data-driven whole-cell strategy,^4^ we constructed a spatial model of a hepatocyte. Briefly, we gathered data for each organelle from a variety of experimental studies such as cryo-electron tomography, live cell fluorescent imaging, and mass spectroscopy. The data is then interpreted to obtain the size, morphology, and number of copies of each hepatocyte organelle. We used set operations and custom-designed codes to faithfully represent the morphology of the organelles.

Hepatocytes are polyhedral-shaped epithelial cells with off-centered nuclei. Therefore, our hepatocyte model has a dimension of 12.3 **µ**m × 12.3 **µ**m × 16 **µ**m,^11^ as shown in Figure 2. The cell organelles that are present in this model include the essential components that are involved in the viral infection process. In total 12 organelles/compartments were included in the model: the plasma membrane, the cytoplasm, the endoplasmic reticulum (ER), the microtubules organizing center (MTOC), the microtubules (MT), the mitochondria, and the nucleus. The nucleus itself is composed of, the nuclear envelope, the nucleoplasm, the nuclear pore complexes (NPCs), the nucleolus, the Cajal bodies, and the nuclear speckles. The three latter components are liquid organelles.^12^

**Figure 2.**
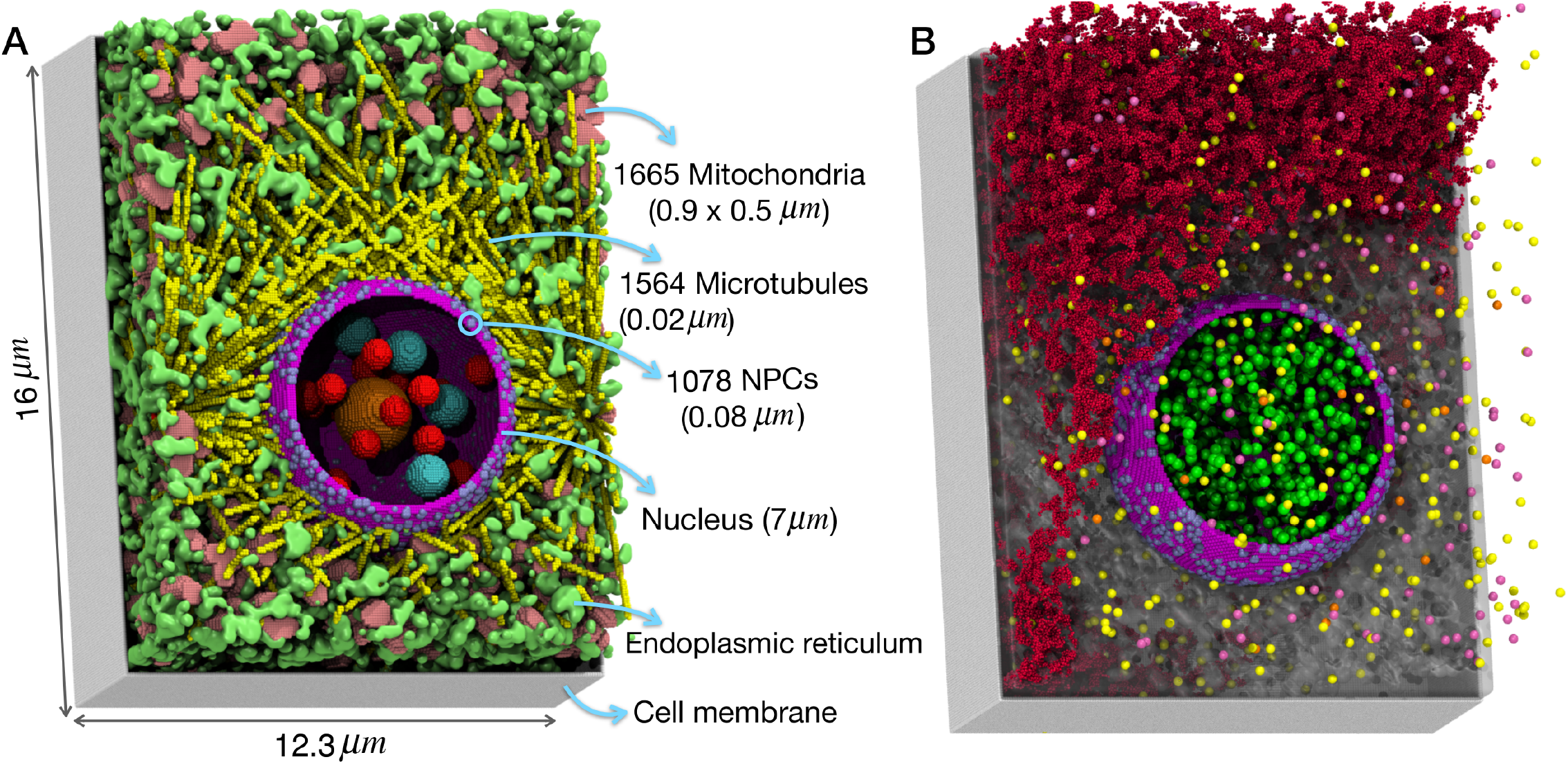
A) The spatial model of the hepatocyte with 12 different organelles and compartments. B) Capsids, shown as yellow (empty), pink (RNA-containing), and orange (mature) are distributed randomly in the cytoplasm which is represented as a shaded grey region. The L proteins are depicted in red at ER (for scene visibility only some of the L proteins are shown). Inside the nucleus, the host cell RNA polymerase II are shown as green spheres.

Similar to our previous HeLa cell model we used to study spliceosomal dynamics,^4^ the ER units were modeled using the cellular automata algorithm, starting from the nuclear envelope and expanding to the plasma membrane, occupying about 7% of the cell volume.^13^ MTOC were modeled as 4 sphero-cylindrical organelles each with the dimension of 0.5 **µ**m × 0.2 **µ**m surrounding the nucleus,^14–16^ from which 1564 MTs with the diameter of 0.064 nm are originated.^17–19^ MTs with random orientation start from the MTOCs and end up close to the cell membrane. MTs can be involved in the transport of viruses in the cytoplasm.^20^ 1665 sphero-cylindrically-shaped mitochondria with a dimension of 0.9 **µ**m × 0.5 **µ**m were randomly distributed in the cytoplasm.^16,21^

The nuclear diameter is 7 **µ**m,^22^ with its envelope covered with 1078 randomly distributed NPCs,^23^ each with a dimension of 0.08 **µ**m,^24^ controlling the traffic of macromolecules between the nucleus and the cytoplasm. The nucleus contains a 0.9 **µ**m-wide nucleolus,^25^ and two distinct liquid organelles: nuclear speckles and Cajal bodies that are both spherically-shaped with radii of 0.35 **µ**m and 0.5 **µ**m, respectively.^12,26^ Hepatocyte organelles are listed in Table 1 along with their dimensions, and the construction of the spatial model of hepatocytes is detailed in Methods.

**Table 1.** Organelles, dimensions and their counts in the model hepatocyte

| Component | Dimension ($\mu\text{m}$) | Number |
| --- | --- | --- |
| Hepatocyte cell size | $16 \times 12.3 \times 12.3$ | 1 |
| Nucleus | 7 (diameter) | 1 |
| mitochondria | $0.9 \times 0.5$ | 1665 |
| MTOC | $0.5 \times 0.2$ | 4 |
| MT | 0.128 (diameter) | 1564 |
| ER | – | 7% cell volume |
| NPCs | 0.08 | 1078 |
| Nucleolus | 1.8 (diameter) | 1 |
| Nuclear speckles | 0.7 (diameter) | 20 |
| Cajal bodies | 1 (diameter) | 4 |

### Kinetic model for the HBV infection

To predict the HBV infection in hepatocytes, we developed a reaction network that can faithfully represent the processes involved in infection. These processes were classified as reactions and diffusion. Because of the small counts of most chemical species involved, we considered the biological events as stochastic processes. Therefore, we use a reaction-diffusion master equation as the theoretical framework that describes the time evolution of the infection (see Methods for details). One of the most important novel aspects of our model is augmenting the reactions to a realistic cellular environment. The spatial degrees of freedom have been shown to be a crucial aspect for infection at different scales.^8,27^ The following biological events were modeled:

0 The viral capsid diffuses in the cytoplasm until reaching an NPC

1. DNA-containing and empty capsids disassemble at NPCs but the RNA-containing capsids are disassembled when they are matured^28^
2. The released relaxed circular DNA (rcDNA) molecules from the DNA-containing capsid at NPCs diffuse in the nucleus until they are converted to covalently closed circular DNA (cccDNA)
3. The cccDNA, the transcriptionally-competent species, is transcribed by the RNA polymerase II and viral messenger RNAs and pgRNA are produced
4. The viral RNA transcripts are transported through the NPCs and are translated in the cytoplasm to produce the viral proteins
5. Viral core proteins assemble to form either empty or genome-containing (pgRNA) capsids
6. The RNA-containing capsid matures
7. Capsids bind to large (L) envelope proteins with correct stoichiometry for envelopment and eventually secretion

The parameters describing the reactions were determined based on available experimental data or modeled based on theoretical approximations. Figure 3 shows our kinetic model that describes the infection life cycle. The design of the reactions together with their associated rates are detailed in Methods. The diffusion coefficients are reported in Table S1.

**Figure 3.**
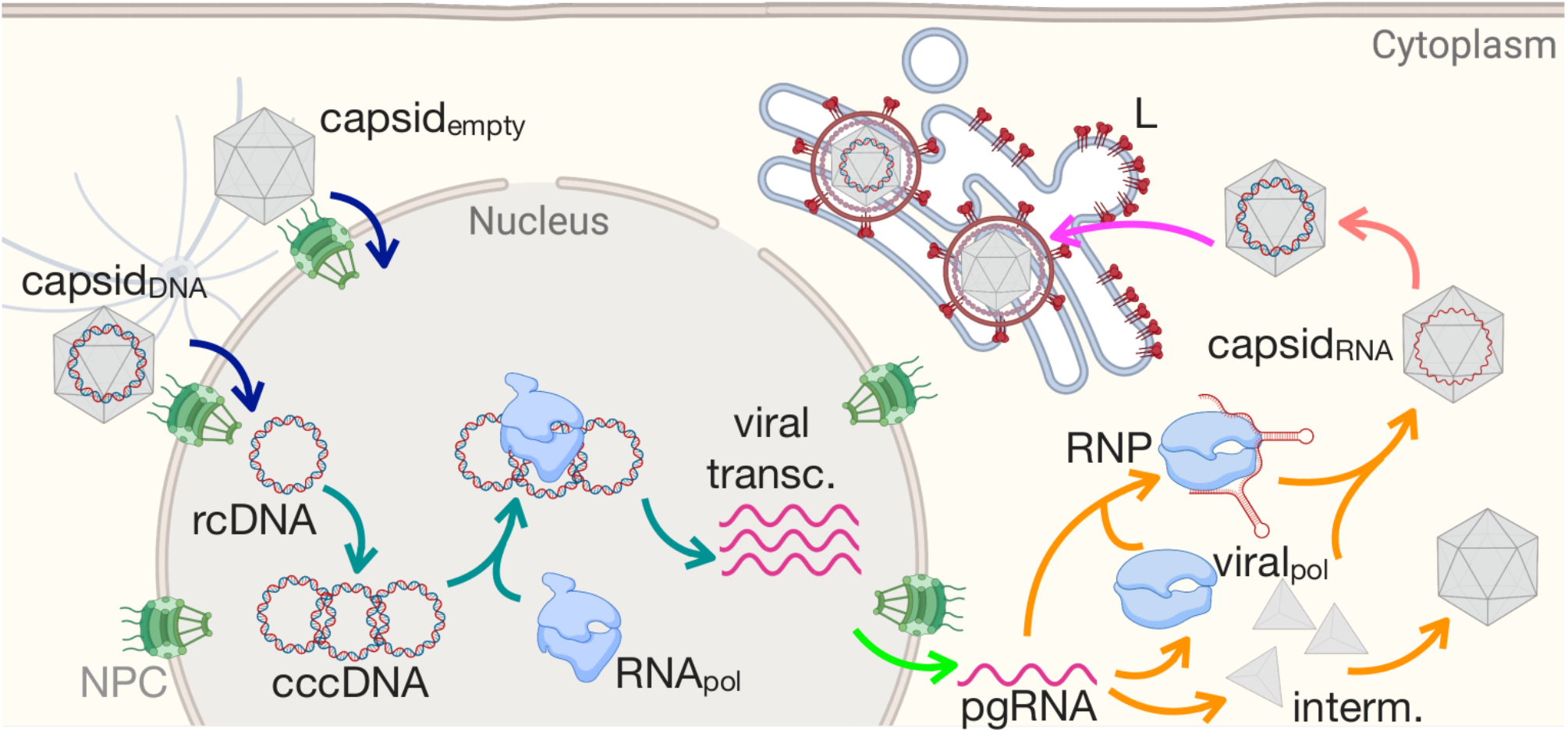
The developed kinetic model describing the HBV infection cycle. The reactions and rates are listed in Table 2.

### Cells switch from the secretion to the amplification pathway in response to envelope protein concentration change

It has been reported that at the start of the infection the amplification pathway is more active, namely capsids are more frequently re-imported to the nucleus, resulting in the enhancement of cccDNA production in the nucleus (Figure 1). In contrast, after the progression of infection, capsids are more frequently enveloped and secreted from the host cell and the secretion pathway becomes more active.^29^ In addition to the cccDNA copy number variation during the infection, one of the factors that is known to affect this pathway switching is the envelope protein copy number, and in particular, large (L) envelope protein that initiates the envelopment.^30^ However, whether in a cell containing an established infection (e.g., with 5 cccDNA, 52 mature capsids, 520 empty capsids^31^) varying L abundance can shift the pathways is not known. This scenario can model the effects of administering the cell with agents that modulate viral gene expression, e.g., short interfering RNA (siRNA) drugs, which may affect the number of L proteins and capsids differentially.^32^

**Table 2.** The reactions added to Table 2 to describe the effects of the drugs on infected cell. Abbreviations are: intermediate core protein complex (interm), cytoplasm (C), and model assumption (M).

| Reaction | Rate | Unit | Compartment | Reference |
| --- | --- | --- | --- | --- |
| interm + interm → capsid<sub>empty</sub>.drug | $1.67 \times 10^{-2}$ | s<sup>-1</sup> | Cytoplasm | <sup>38</sup>, M. |
| Capsid<sub>DNA</sub>.drug → 0 | $1.67 \times 10^{-3}$ | s<sup>-1</sup> | Cytoplasm | <sup>38</sup> |

Starting from a value of 5×10^5^ copies of L protein,^10^ we simulated our developed infection cycle of HBV, incorporated in the entire spatial model of a hepatocyte for 10 minutes of biological time with varying L abundance (see Methods for details). We monitored different viral species as a function of time to determine how infection is evolving. Most importantly, we tracked the hallmark of chronic infection, i.e., cccDNA molecules inside the nucleus at the end of our simulations. As we decrease the L abundance, the cccDNA copy number increases, indicating that more capsids are re-imported to the nucleus, disassembled at NPCs to release their viral DNA into the nucleus, and hence, the amplification pathway activates as shown in Figure 4A. In contrast, for high L abundance the counts of cccDNA remain close to its starting value, indicating that capsids are less re-imported to the nucleus and the secretion pathway is more active. Therefore, the reduction in L counts (or increase in siRNA concentration), results in switching between pathways from secretion to amplification.

**Figure 4.**
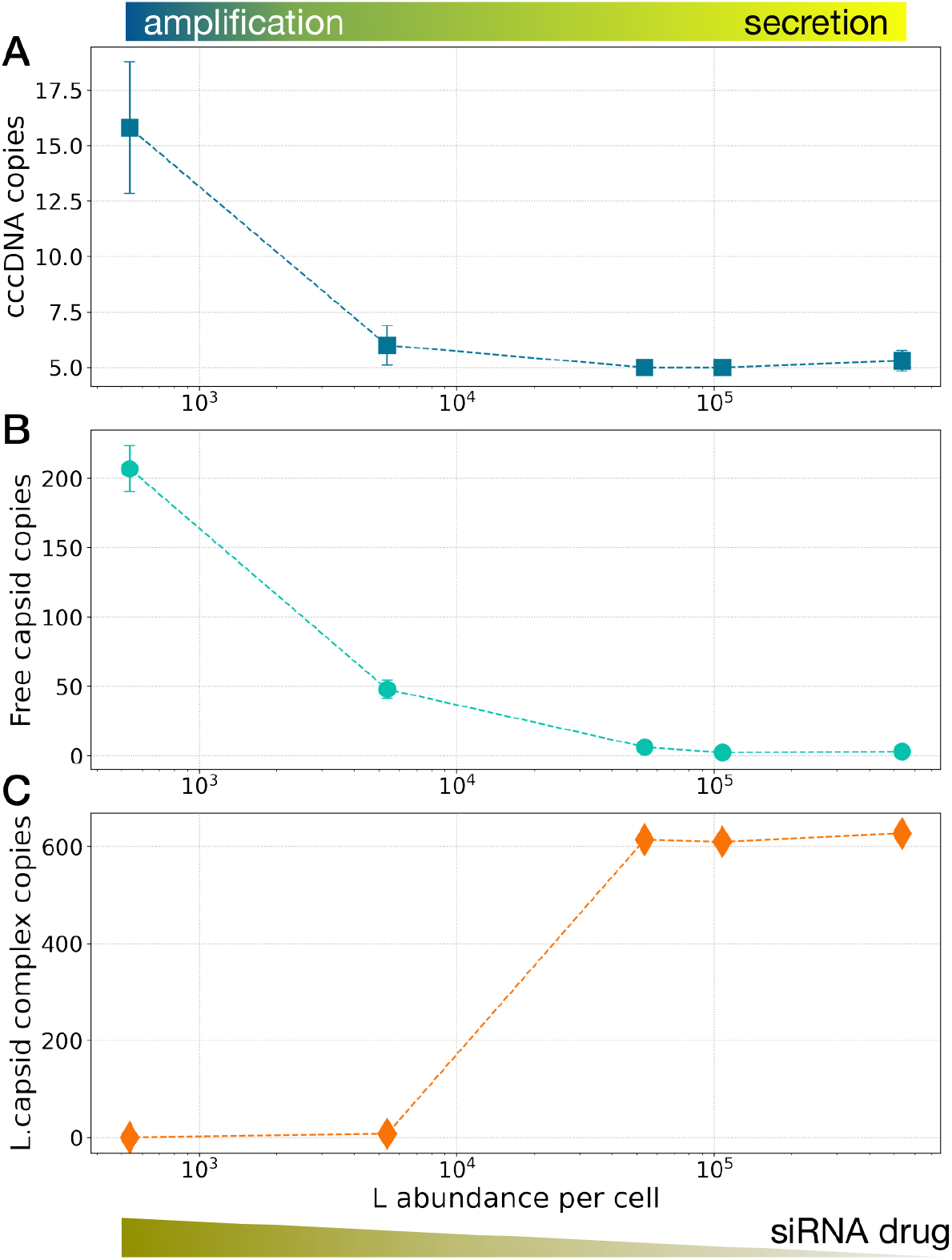
Envelope protein (L) or as a proxy for siRNA drugs, can switch the dominant replication pathway from secretion to amplification resulting in increased A) cccDNA copy number, B) available capsids (both mature and empty), and C) capsids in complex with L proteins. The error bars show the standard deviations calculated from 10 simulation replicas.

As the active pathway switches to amplification, capsids (mature and empty) are less frequently secreted, and therefore the number of cytoplasmic capsids is increased (Figure 4B). Consistently, number of the L proteins bound to the capsid at ER (L.capsid complex), which are ready to continue with the rest of envelopment process and ultimately secretion, decreases (Figure 4C). The results are independent of the initial concentrations of different species pointing to their universality. These results relate L protein and cccDNA concentration quantitatively and demonstrate that our whole-cell model is responsive to perturbations, and capable of reproducing expected behavior. More interestingly, they show that this class of drugs can lead to a contained, but increased infection within the host cell because L protein downregulation leads to more viral DNA in the nucleus. Thus, development of agents to modify viral gene expression including siRNAs and viral transcriptional inhibitors needs to take into consideration of the relative effects on the capsids and envelope proteins and to be pursued in combination with other drugs to reduce viral DNA such as RT inhibitors.^33^

### Different capsid disassembly mechanisms lead to differential infection outcome

We have shown that mutations at specific regions of the core (capsid) protein can increase the level of cccDNA production with respect to the wild type capsids.^34,35^ Additionally, we predicted the presence of disassembly hotspots (weak points) on the capsid,^36^ and showed that core protein mutations can destabilize the capsid, thus enhancing its disassembly.^37^

We hypothesize that capsid mutants that increase cccDNA production destabilize the capsid only enough to accelerate its disassembly at NPCs but prevents its cracking in the cytoplasm. To test this hypothesis, we simulated the effects of the capsid mutants by varying the capsid disassembly time, as a measure of different degrees of capsid destabilization. We started from capsid disassembly time of ∼10 minutes, determined by *in vitro* time-resolved small angle X-ray scattering experiments,^38^ and changed it in the range of 1 to 100 minutes, all the disassembly events occurs at NPCs. Similar to above, these simulations incorporated the entire infection cycle in the realistic model of a hepatocyte. Figure 5A shows a snapshot of the simulation at which capsids (orange and yellow spheres) diffused and bound to the NPCs. By decreasing the disassembly time, the normalized counts of capsids at NPCs (i.e., the ratio of the total number of capsids bound to NPCs at the end of the simulation to the same quantity after 30 s of simulation time), decreases (Figure 5B). This result indicates that due to the faster disassembly of the mutated capsids, fewer capsids remain bound to NPCs. Following the accelerated disassembly for mutant capsids, the number of synthesized cccDNA molecules increases, which leads to enhanced infection (Figure 5C). If on the other hand, the capsid is hyper-destabilized and the disassembly occurs in the cytoplasm (Figure 5C, orange dot), no cccDNA is replicated beyond its initial value (5), which is obviously the preferred outcome to cccDNA increase for therapy, which we discuss next.

**Figure 5.**
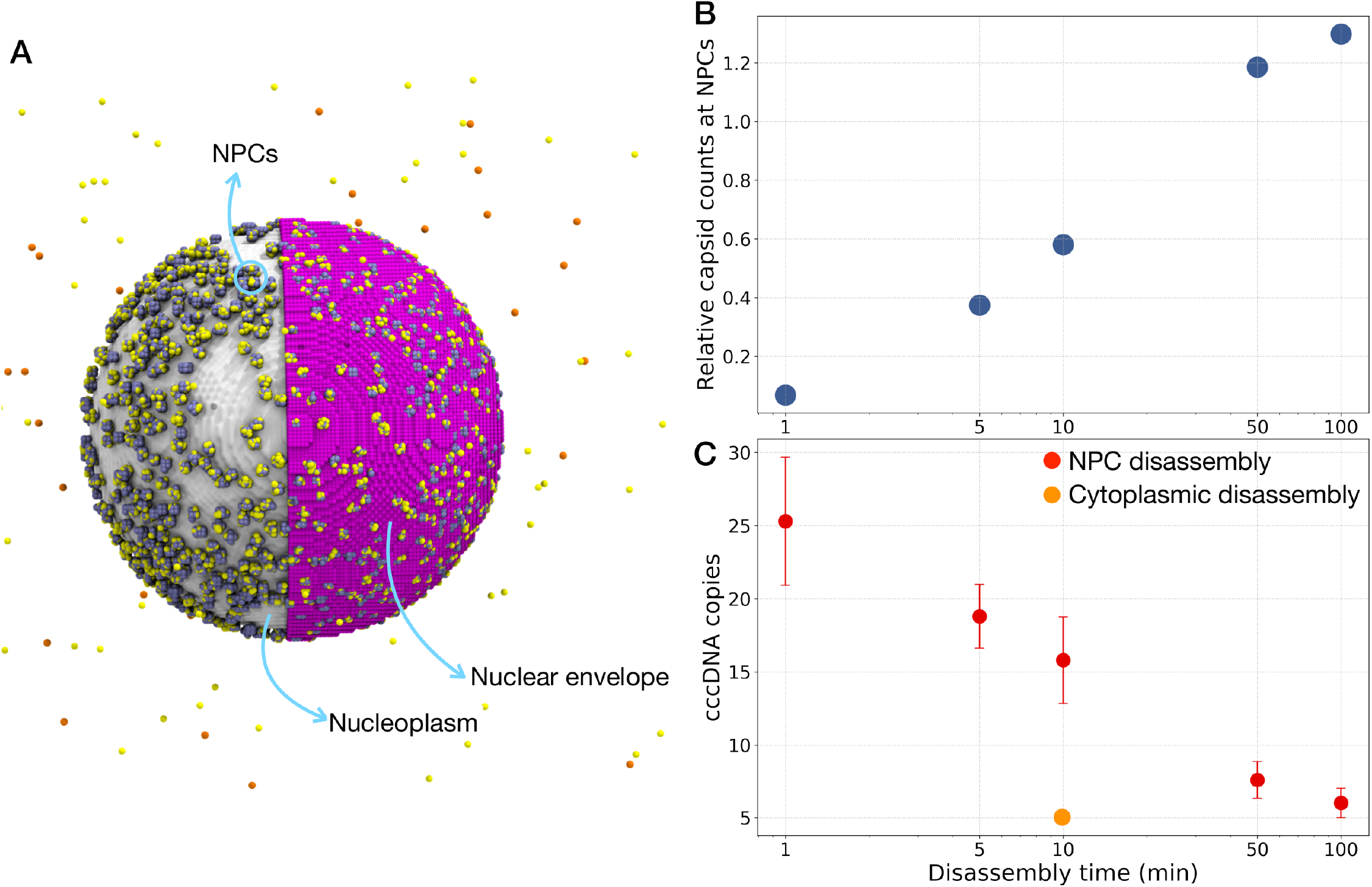
Effect of capsid stability on interaction with NPCs and cccDNA counts. (A) Snapshot of capsids (shown in yellow and orange) in cytoplasm and bound to NPCs (colored in iceblue). Half of the nuclear envelope is shown in purple. By decreasing the disassembly time: B) the relative number of capsids in NPCs (namely, the ratio of the total number of capsids bound to NPCs at the end of the simulation to the same quantity after 30 s of simulation time) decreases, and C) the number of cccDNA molecules increases provided that the mutant capsids remain stable enough not to crack in the cytoplasm. The error bars show the standard deviations calculated from 10 simulation replicas.

### Simulations predict an effective drug concentration and the best timing for drug administration

HBV capsid inhibitor (e.g. heteroaryldihydropyrimidines, and sulfamoylbenzamides) can accelerate capsid assembly and by destabilizing the capsid, can lead to its enhanced disassembly.^39^

It was shown that these drugs can reduce the cccDNA copy number during a *de novo* infection (when the infection is being initiated at a host cell) but unconstructively they increase the cccDNA copy number during the amplification, which defeats their purpose.^39^ This is possibly because these compounds may induce capsid disassembly at the NPC, similar to the effects of capsid mutations that enhance cccDNA formation via increased destabilization of mature capsids, as described above. We hypothesize that, if drug binding induces capsid disassembly to occur in the cytoplasm, the capsid inhibitors can indeed decrease the number of cccDNA in the nucleus, turning them into more effective drugs.

We use our whole-cell computational platform to test this hypothesis by adding additional reactions to the kinetic model to simulate the effects of drugs acting on an infected cell. As mentioned above, drugs binding to core proteins accelerates capsid assembly. Thus, we assumed that the capsid assembly time is reduced by a factor of 10 from its experimentally determined value (Table 2).^38^ Additionally, the initial condition for this simulation set is that drugs are bound to a subset of capsids (capsid_DNA_.drug). The drug-bound capsids then disassemble in the cytoplasm and in contrary to the drug-free simulations, the rcDNA is degraded in the cytoplasm and therefore, nothing is produced. The additional reactions to account for drugs mechanism of action are listed in Table 3.

To determine an effective concentration for the drug, we varied the fraction of capsids that are drug bound. Figure 6A shows that 20% drug-bound capsids, are not enough to significantly reduce the cccDNA copy number. Drug dosage needs to be increased to at least bind to 50% of capsids, and ultimately reaching a factor of 2 reduction in cccDNA copy number when increasing the drug concentration to bind to 80% of capsids.

**Figure 6.**
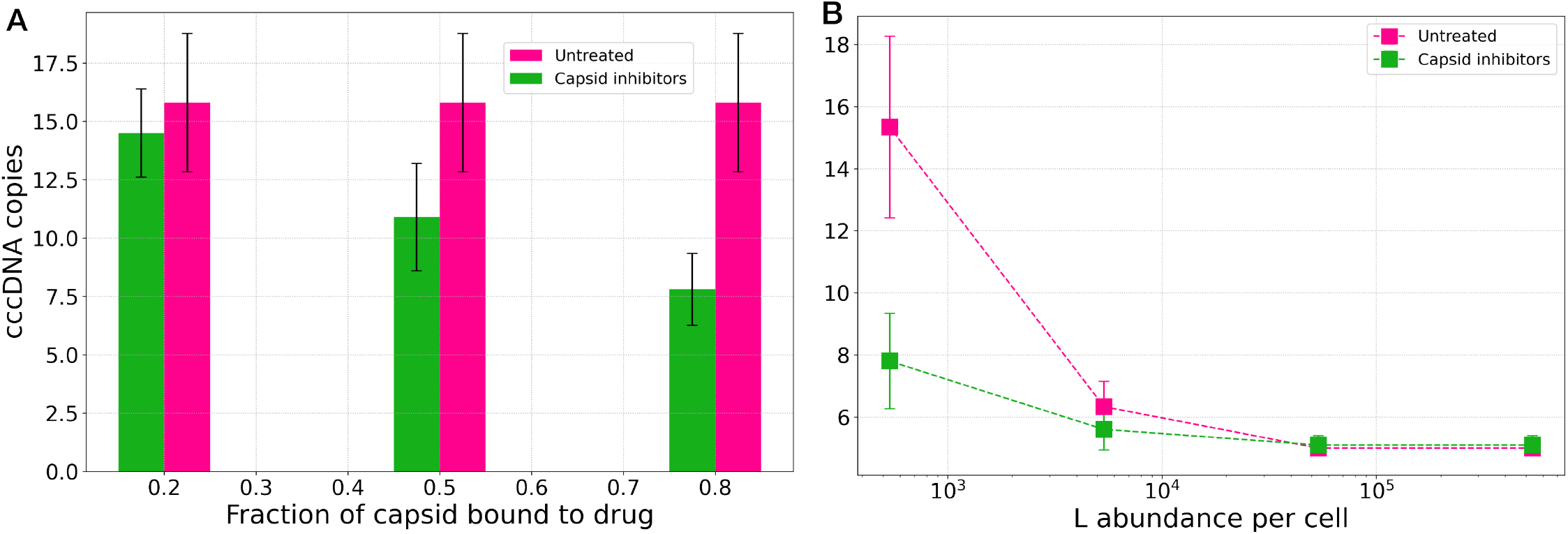
Efficacy of a hypothetical capsid inhibitor drug treatment that promotes cytoplasmic dissociation of the capsid. (A) The effect of drug concentration on cccDNA copies when treated (green) as compared to and untreated (magenta, for reference) cells. (B) The timing of drug administration matters: drugs are only effective at the start of infection, or when L abundance is low. The error bars show the standard deviations calculated from 10 simulation replicas.

Can the timing of drug administration change the effectiveness of the drug? To answer this question, we simulated the effect of capsid inhibitors in different stages of infection. Specifically, we varied L protein abundance to generate host cells in amplification-active vs. secretion-active pathways (Figure 4). The results in Figure 6B show that at the start of infection when amplification pathway is active and the L protein count is low, drug treatment effectively reduces the cccDNA copy number, which is the hallmark of infection. But as the infection progresses, and the secretion pathway becomes more active or at high L counts, even drugs bound to 80% of capsids will not reduce the number of cccDNA molecules in the nucleus. This is because when secretion is dominant, capsids mostly are enveloped and secreted, leaving small number of apo capsids (not bound to drugs) that can be re-imported to the nucleus.

Therefore, our results suggest that for capsid inhibitors to be effective in decreasing the cccDNA copy number during the amplification pathway, the drug-bound capsids: 1) need to be disassembled in the cytoplasm; and 2) the drug concentration should be high enough to bind to at least 50% of capsids.

## Discussion and conclusions

We used a data-driven approach to construct a whole-cell computational platform for investigating the viral infection cycle, virus-host cell interactions and infection-drug interactions. Our focus in this study was on HBV, that World Health Organization aims to eliminate by the year 2030. We developed a spatially-resolved model for a hepatocyte (liver cell), the host cell of HBV, and a kinetic model, which is a series of reactions that describe the viral infection inside the cell.

We simulated our whole-cell platform stochastically for 10 minutes of biological time. We found that, with an established infection, decreasing the abundance of viral envelope (L) proteins can switch the infection pathway from dominantly secreting the capsids to re-importing them back to the nucleus, resulting in cccDNA amplification. The L protein concentration, relative to the capsid, can be a proxy for a class of drugs modulating viral gene expression (e.g., siRNA), indicating that these drugs can lead to an increase in viral cccDNA, counterproductive to the therapeutic goal. Our simulations showed that the degree to which capsid mutants destabilize capsid structure can lead to differential outcomes for cccDNA copy number: severe destabilization which leads to capsid cracking at the cytoplasm does not affect the cccDNA copy number, in contrast, a milder destabilization, still keeping the disassembly at NPCs, increases the cccDNA copy number. Thus, drugs designed to affect capsid stability must carefully balance these two scenarios. Finally, by simulating the effects of capsid inhibitor drugs, we predicted the dosage and timing of drug administration that can be most effective for reducing the cccDNA copy number.

Our model could be used to test a variety of scenarios of variable capsid stability, depending on the mode of action of a drug. The binding affinity to nuclear pore machinery could be reduced by a core protein mutation without compromising capsid stability. Capsid stability could be enhanced so disassembly at the nuclear pore is inefficient, and the question is how much stabilization would be effective? And as noted above, disassembly could be promoted in the cytoplasm, in which case the question is to what extent undissociated capsids in equilibrium with the disassembled capsid can still lead to breakthrough of an infection. The model can probe what ranges of destabilization or reduced binding to the pore complex would be needed to mitigate infection effectively, providing goal posts for drug design, including multimodal design where all of these effects can interact in real hepatocytes and in our model.

Our computational platform can be scaled to incorporate other host cell processes or viral pathways and simulate other HBV drugs as well as combination therapies. The platform enables researchers to identify the most crucial viral pathways whose targeting with existing or newly-designed drugs can yield the most effective response. Finally, the platform is extendable to other viruses by changing the kinetic model and if necessary, by modifying the spatial organization of the cell.

## Methods

### Construction of a spatially-resolved hepatocyte

To accurately represent HBV infection which involves multiple cellular compartments, we constructed a hepatocyte with spatial representation. We used a variety of experimental data describing e.g., the size, morphology, relative mass fraction of specific components to inform model cell construction. To build the cell, we used a constructive solid geometry (CSG) approach, that utilizes set operations (e.g., unions, differences, intersections) to combine basic geometric objects are combined. Since LM requires that each location within the space be defined as a single site-type the various CSG objects were stenciled onto the simulation lattice in “depth order” (also called the “Painter’s algorithm”).^3,40,41^ Overall, our model consists of 13 different site-types including: 1) extracellular space, 2) plasma membrane, 3) cytoplasm, 4) mitochondria, 5) endoplasmic reticulum, 6) microtubules, 7) microtubules organizing center, 8) nuclear envelope, 9) nucleoplasm, 10) Cajal bodies, 11) nuclear speckles, 12) nucleolus, and 13) nuclear pore complexes. Because of the cuboidal shape of hepatocytes, the simulation volume was constructed as a cubic box with a dimension of 16.38 μm side-length. The space was discretized into cubic lattices with a size of 64 nm, within and among which reactions and diffusions occur. In addition to the cell dimension of ∼15 μm, the volume of a hepatocyte measured as 6.9 pL.^42^ To reduce the computational cost of our simulations we chose the former dimension. Plasma and nuclear membranes were implemented as a thin sheet of lattice points (128 nm thick) separating the extracellular space, the cytoplasm and the nucleus. Mitochondria were modeled as randomly oriented spherocylinders placed within the cytoplasm. We have shown previously that the network-like morphology of the mitochondria does not change the cytoplasmic diffusions and reaction of species significantly.^4^ Following successful generation of ER for HeLa cells using the game of life algorithm,^4^ we used a similar algorithm for a randomized ER generation in the cytoplasm. Four MTOCs oriented randomly but all within the same distance of ∼2 μm from the nuclear envelope, are the starting points of MTs, with a thickness of 2 lattice sites (128 nm). Nuclear pore complexes (NPCs) were embedded in the nuclear envelope. NPCs were constructed as a set of spheres of radii 0.08 μm to ensure connectivity from the nucleus to the cytoplasm. Inside the nucleus, similar to HeLa cells, nuclear speckles and Cajal bodies were modeled as spheres placed randomly. In addition, we added a spherical nucleolus with a diameter of 1.8 μm to the nucleus of hepatocyte. A list of organelles and their dimensions can be found in Table 1.

Our aim was to make this cell model more complete than our previously developed HeLa cell and to include all the organelles that are involved in viral infection.^4^ However, the organelles that contribute less than 1% to the total cell composition,^13^ such as endosomes, lysosomes, and peroxisomes, as well as chromatin have not been included in the current hepatocyte model.

### Kinetic model development and parameter estimation

Another important part of our computational platform is to faithfully represent the biological events through a series of reactions and diffusion of different species. The details of different infection steps presented in Figure 3 are described below. The reactions describing the degradation of different species that were included in the kinetic model are presented in Table S2.

#### 1 Capsid disassembly at NPCs

when the DNA-containing capsid reaches the nucleus to deliver the viral genome, the capsid is thought to disassemble at NPCs. The rate of these reactions was determined by *in vitro* experiments to be 1.67×10^−3^ s^−1^.^38^

| Step 1 reactions | Rate | Unit | Compartment | Reference |
| --- | --- | --- | --- | --- |
| $\text{capsid}_{\text{DNA}} \rightarrow \text{rcDNA}$ | $1.67 \times 10^{-3}$ | $\text{s}^{-1}$ | NPC | 38 |
| $\text{capsid}_{\text{empty}} \rightarrow 0$ | $1.67 \times 10^{-3}$ | $\text{s}^{-1}$ | NPC | 38 |

#### 2 rcDNA repair and formation of cccDNA

rcDNA is repaired and converted to covalently-closed circular DNA (cccDNA) which is the transcriptionally competent genome species.^7^ The rate of this reaction was based on the kinetics of the repair reactions for the plus- and negative-strands, studied in biochemical repair systems.^43^

| Step 2 reaction | Rate | Unit | Compartment | Reference |
| --- | --- | --- | --- | --- |
| $\text{rcDNA} \rightarrow \text{cccDNA}$ | $3.3 \times 10^{-3}$ | $\text{s}^{-1}$ | Nucleus | 43 |

#### 3 cccDNA transcription at nucleus

The host cell RNA polymerase II (RNApol) then binds to the cccDNA. The binding/unbinding rates of this reaction were approximated from.^44^ With a transcription rate of 50 nucleotide/s,^45^ and the genome length for pgRNA, preC, X, M, S, and L proteins of 2.5 Kbase (b), 3.5 Kb, 0.7 Kb, 2.1 Kb, 2.1 Kb, and 2.4 Kb, respectively,^46^ we calculated the rate of each reaction.

| Step 3 reactions | Rate | Unit | Compartment | Reference |
| --- | --- | --- | --- | --- |
| $\text{cccDNA} + \text{RNA}_{\text{pol}} \rightarrow \text{cccDNA}.\text{RNA}_{\text{pol}}$ | $0.33 \times 10^6$ | $\text{M}^{-1} \text{s}^{-1}$ | Nucleus | 44 |
| $\text{cccDNA}.\text{RNA}_{\text{pol}} \rightarrow \text{cccDNA} + \text{RNA}_{\text{pol}}$ | $1 \times 10^{-3}$ | $\text{s}^{-1}$ | Nucleus | 44 |
| $\text{cccDNA}.\text{RNA}_{\text{pol}} \rightarrow \text{cccDNA} + \text{RNA}_{\text{pol}} + \text{pgRNA}$ | 0.02 | $\text{s}^{-1}$ | Nucleus | 45,46 |
| $\text{cccDNA}.\text{RNA}_{\text{pol}} \rightarrow \text{cccDNA} + \text{RNA}_{\text{pol}} + \text{LSmRNA}$ | 0.0208 | $\text{s}^{-1}$ | Nucleus | 45,46 |
| $\text{cccDNA}.\text{RNA}_{\text{pol}} \rightarrow \text{cccDNA} + \text{RNA}_{\text{pol}} + \text{SmRNA}$ | 0.0238 | $\text{s}^{-1}$ | Nucleus | 45,46 |
| $\text{cccDNA}.\text{RNA}_{\text{pol}} \rightarrow \text{cccDNA} + \text{RNA}_{\text{pol}} + \text{XmRNA}$ | 0.0714 | $\text{s}^{-1}$ | Nucleus | 45,46 |
| $\text{cccDNA}.\text{RNA}_{\text{pol}} \rightarrow \text{cccDNA} + \text{RNA}_{\text{pol}} + \text{preCmRNA}$ | 0.014 | $\text{s}^{-1}$ | Nucleus | 45,46 |

#### 4 Viral proteins translation

After the transport of RNA transcripts from the nucleus to the cytoplasm through NPCs, in our model, they implicitly bind to the ribosomes and translation occurs. To optimize the computational cost of our simulations, we introduced lumped reactions for core protein monomer and dimer species. Namely, when pgRNA is translated, an intermediate species (denoted as “interm” below) composed of 60 core protein dimers is directly produced. PgRNA translation also produces a viral polymerase (viral_pol_). The rates of all translation reactions were calculated based on translation speed of 11 amino acids/s^47^ and their known number of nucleotides in mRNA transcripts.^46^

| Step 4 reactions | Rate | Unit | Compartment | Reference |
| --- | --- | --- | --- | --- |
| $\text{pgRNA} \rightarrow \text{interm} + \text{pgRNA}$ | 0.0131 | $\text{s}^{-1}$ | Cytoplasm | 47 |
| $\text{pgRNA} \rightarrow \text{viral}_{\text{pol}} + \text{pgRNA}$ | 0.0131 | $\text{s}^{-1}$ | Cytoplasm | 47 |
| $\text{LSmRNA} \rightarrow \text{L} + \text{LSmRNA}$ | 0.0139 | $\text{s}^{-1}$ | Cytoplasm | 47 |
| $\text{SmRNA} \rightarrow \text{M} + \text{SmRNA}$ | 0.0156 | $\text{s}^{-1}$ | Cytoplasm | 47 |
| $\text{SmRNA} \rightarrow \text{S} + \text{SmRNA}$ | 0.0156 | $\text{s}^{-1}$ | Cytoplasm | 47 |
| $\text{XmRNA} \rightarrow \text{X} + \text{XmRNA}$ | 0.048 | $\text{s}^{-1}$ | Cytoplasm | 47 |
| $\text{preCmRNA} \rightarrow \text{preCmRNA} + \text{preC}$ | 0.0132 | $\text{s}^{-1}$ | Cytoplasm | 47 |

#### 5 Capsids assembly

A viral capsid is then assembled by first the binding of pgRNA and viral polymerase to form the ribonucleoprotein (RNP) complex. The rates of these reactions were approximated to the binding of cccDNA to RNA_pol_.^44^ Then in two steps RNP binds to the intermediate core protein species and form RNA-containing capsid (capsid_RNA_). For the empty capsid (capsid_empty_) assembly instead, two intermediate species come together. The rates of these reactions were based on *in vitro* experimental measurements.^38^ Because empty capsids are ca. 5-10 times more abundance that the RNA-containing capsids inside the cell,^48^ the rate of the assembly of RNA-containing capsids was multiplied by a factor of 0.2. Note that because the experimental unit for capsid assembly measurement was in second, for the assembly reactions the stochastic rate was directly mentioned in the reactions table.

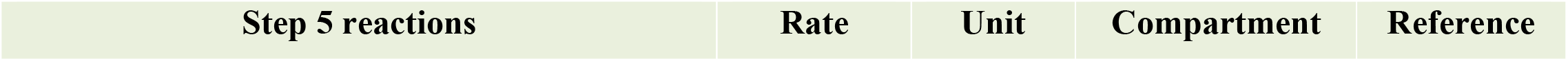
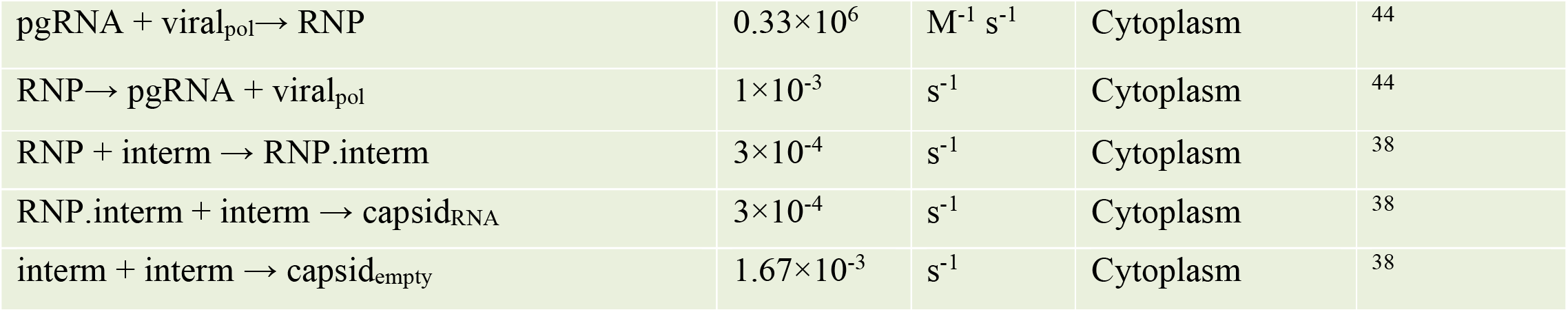

| Step 5 reactions | Rate | Unit | Compartment | Reference |
| --- | --- | --- | --- | --- |
| pgRNA + viral<sub>pol</sub> → RNP | 0.33×10<sup>6</sup> | M<sup>-1</sup> s<sup>-1</sup> | Cytoplasm | 44 |
| RNP → pgRNA + viral<sub>pol</sub> | 1×10<sup>-3</sup> | s<sup>-1</sup> | Cytoplasm | 44 |
| RNP + interm → RNP.interm | 3×10<sup>-4</sup> | s<sup>-1</sup> | Cytoplasm | 38 |
| RNP.interm + interm → capsid<sub>RNA</sub> | 3×10<sup>-4</sup> | s<sup>-1</sup> | Cytoplasm | 38 |
| interm + interm → capsid<sub>empty</sub> | 1.67×10<sup>-3</sup> | s<sup>-1</sup> | Cytoplasm | 38 |

#### 6 Reverse transcription of the viral RNA and maturation of capsid

An important step in the life cycle of HBV is the reverse transcription of viral RNA inside the capsid and formation of DNA-containing capsids, following by capsid maturation processes. This process is known to be slow with a half-life of 12 hours.^10^

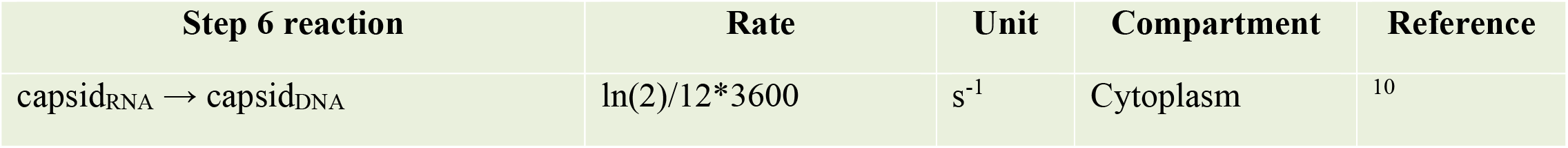

| Step 6 reaction | Rate | Unit | Compartment | Reference |
| --- | --- | --- | --- | --- |
| capsid<sub>RNA</sub> → capsid<sub>DNA</sub> | ln(2)/12*3600 | s<sup>-1</sup> | Cytoplasm | 10 |

#### 7 Capsid binding to envelope proteins

All three types of capsids can acquire an envelope and get secreted from the cell, but with different ratios.^7^ The viral envelope consists of three types of proteins of small, medium and large, embedded in lipid membrane^49^ with the stoichiometry of 80, 20, and 20, respectively.^50^ The envelopment is initiated by binding of the viral capsid to large (L) proteins that are localized at ER.^49^ We explicitly simulated the binding of the capsid to each of the 20 L proteins. The binding rate constant for L protein binding to a mature capsid (K_bm_) was estimated as a diffusion-limited reaction (D.L. in table below), and the unbinding rate constant was calculated (C.) from the measured dissociation constant of the inhibitory peptides binding to a capsid.^51^ Inside the cell the abundance of capsid_RNA_ is at least a factor of 2 higher than mature capsids.^52,53^ In contrary, the abundance of the RNA-containing capsids that are secreted from the cell is 1000 times lower than mature capsids.^7^ Therefore, the binding rate constant of the capsid_RNA_ binding to L proteins, was estimated by scaling K_bm_ by a factor of 5×10^−4^. Similarly, based on the difference of the ratio of empty capsids that are secreted from the cell and their intracellular abundance, the binding rate constant of the capsid_empty_ binding to L proteins was scaled by a factor of 10.

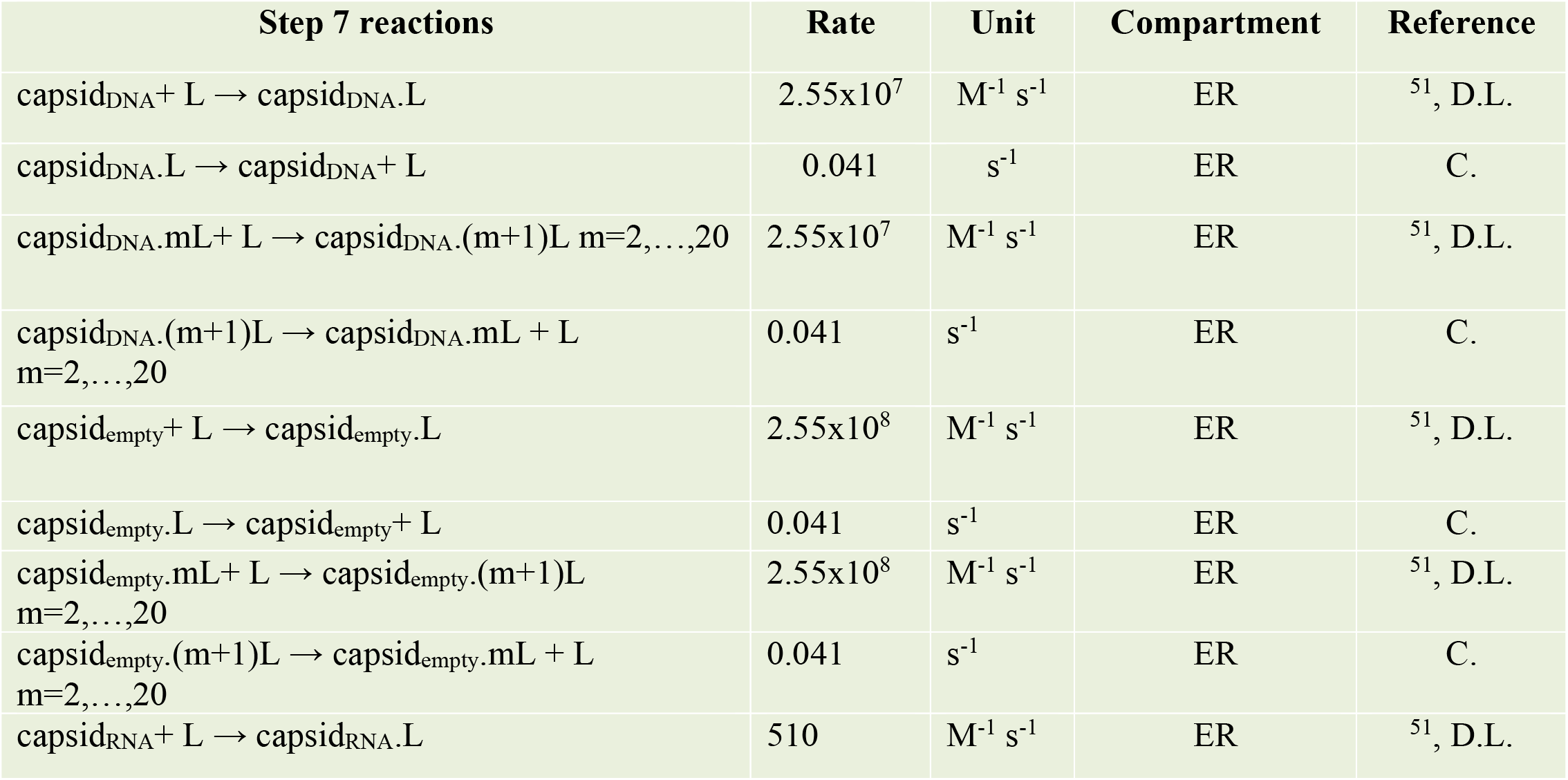
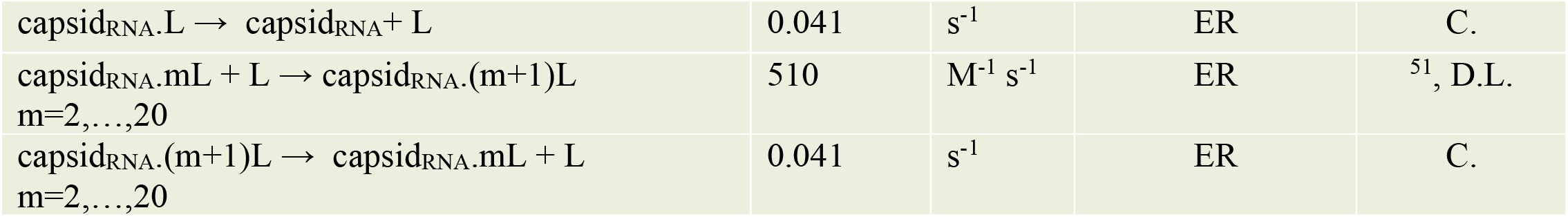

| Step 7 reactions | Rate | Unit | Compartment | Reference |
| --- | --- | --- | --- | --- |
| capsid<sub>DNA</sub> + L → capsid<sub>DNA</sub>.L | 2.55×10<sup>7</sup> | M<sup>-1</sup> s<sup>-1</sup> | ER | <sup>51</sup>, D.L. |
| capsid<sub>DNA</sub>.L → capsid<sub>DNA</sub> + L | 0.041 | s<sup>-1</sup> | ER | C. |
| capsid<sub>DNA</sub>.mL + L → capsid<sub>DNA</sub>.(m+1)L m=2,...,20 | 2.55×10<sup>7</sup> | M<sup>-1</sup> s<sup>-1</sup> | ER | <sup>51</sup>, D.L. |
| capsid<sub>DNA</sub>.(m+1)L → capsid<sub>DNA</sub>.mL + L m=2,...,20 | 0.041 | s<sup>-1</sup> | ER | C. |
| capsid<sub>empty</sub> + L → capsid<sub>empty</sub>.L | 2.55×10<sup>8</sup> | M<sup>-1</sup> s<sup>-1</sup> | ER | <sup>51</sup>, D.L. |
| capsid<sub>empty</sub>.L → capsid<sub>empty</sub> + L | 0.041 | s<sup>-1</sup> | ER | C. |
| capsid<sub>empty</sub>.mL + L → capsid<sub>empty</sub>.(m+1)L m=2,...,20 | 2.55×10<sup>8</sup> | M<sup>-1</sup> s<sup>-1</sup> | ER | <sup>51</sup>, D.L. |
| capsid<sub>empty</sub>.(m+1)L → capsid<sub>empty</sub>.mL + L m=2,...,20 | 0.041 | s<sup>-1</sup> | ER | C. |
| capsid<sub>RNA</sub> + L → capsid<sub>RNA</sub>.L | 510 | M<sup>-1</sup> s<sup>-1</sup> | ER | <sup>51</sup>, D.L. |
| $\text{capsid}_{\text{RNA}}.\text{L} \rightarrow \text{capsid}_{\text{RNA}} + \text{L}$ | 0.041 | $\text{s}^{-1}$ | ER | C. |
| $\text{capsid}_{\text{RNA}}.\text{mL} + \text{L} \rightarrow \text{capsid}_{\text{RNA}}.(m+1)\text{L}$<br>$m=2,\dots,20$ | 510 | $\text{M}^{-1} \text{s}^{-1}$ | ER | <sup>51</sup>, D.L. |
| $\text{capsid}_{\text{RNA}}.(m+1)\text{L} \rightarrow \text{capsid}_{\text{RNA}}.\text{mL} + \text{L}$<br>$m=2,\dots,20$ | 0.041 | $\text{s}^{-1}$ | ER | C. |

### RDME simulations methodology in LM

Following detailed discussion in refs.^4,40,54^ in details, we use formalism known as the reaction-diffusion master equation (RDME), which describes the state of a stochastic system to be in a specified spatial and compositional state, for the simulations in this paper. The “state” for our systems of interest, comprises of the number of each interacting species along with their positions. In RDME simulations using LM, the geometry of the system is first discretized to a cubic grid with a lattice spacing λ. Additionally, a fundamental time step τ is specified. The time evolution of the probability of the system to be in specific state **x**, containing number of molecules for each of the *N*_*sp*_ species present at each subvolume υ∈V, is determined by the rates of change due to reaction and diffusion, defined by operators **R** and **D**, respectively:

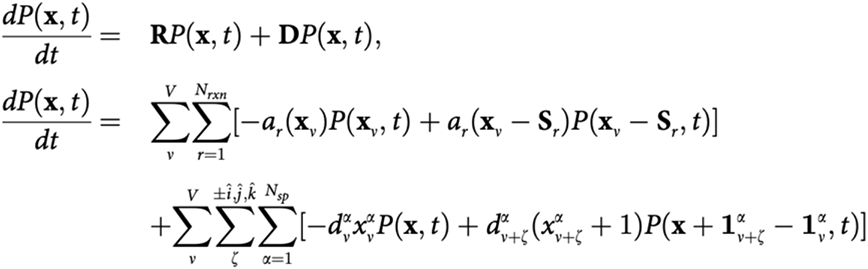

The first two terms of this equation define the probability flux within each subvolume due to reactions: *a*_*r*_(**xυ**) is the reaction propensity for reaction *r* of *N*_*rxn*_ reactions and **S***r* is the *N* × *R* stoichiometric matrix describing the net change in molecules number when a reaction occurs. The last two terms describe the rate of the change of the probability due to molecules’ propensity to diffuse between neighboring subvolumes. *d*_υα_ is the diffusive propensity for a molecule of species *α* to jump to a neighboring subvolume, and is related to the macroscopic diffusion. The first part of the diffusion operator defines the probability flux out of the current state due to the diffusion of molecules from subvolume υ; and the second part of the diffusion operator, describes the diffusion of molecules into the current state from subvolumes, υ+η, where η defines the neighboring subvolume in ±*x*, ±*y*, ±*z* directions. Using an approximation for sampling the RDME equation, as implemented in Lattice Microbes software, simulations of biologically relevant timescales (10 mins) for the hepatocyte model on multiple GPU hardware became possible.

### RDME simulation details

the goal of our simulations was to study how a specific infection parameter/drug affects the infection. Therefore, all our simulations started from an infected cell with an steady-state viral components that were determined previousely.^10,31^ All simulations started with 5 cccDNA, 52 mature capsids, 260 RNA-containing capsids, 520 empty capsids, 59 pgRNA, and 680 RNA polymerase II, and ∼ 500 L proteins when its counts is not specified. Our whole-cell model contains a total of 169 reactions occurring between 89 chemical species. All simulation time steps were 9.3 × 10^−5^ s, and lattice spacing was 64 nm. The chemical species counts were written every 10 time step. All simulations were performed for 10 minutes of biological times on multi-GPU clusters. For each simulation condition we generated 10 replicas and calculated the mean and standard deviation over the replicas.

## Acknowledgments

M.G. was supported by the James R. Eiszner Chair. Z.G. was supported by the Department of Chemistry at the University of Illinois at Urbana-Champaign, as well as NSF MCB 2205665 (Gruebele) and NIH P41-104601 (Tajkhorshid). E.T. was supported by NIH P41-104601. J.H. was supported by R37 AI043453. O.N. was supported by the Carle-Illinois College of Medicine as part of a Discovery Learning research project. The computational resources were provided by the NIH Resource P41-104601.

