## Supplementary material for "A Spatial Whole-Cell Model for Hepatitis B Viral Infection and Drug Interactions": Table S1 and Table S2

Table S1. Diffusion coefficients of species involved in the reactions

| Species | Diffusion coeff. | Compartment | Reference |
|---|---|---|---|
| $capsid_{DNA}$ | $7.4\times10^{-12}$ | Cytoplasm/ER | [1] |
| $capsid_{DNA}$ | $16\times10^{-12}$ | NPC | [1] |
| $capsid_{empty}$ | $7.4\times10^{-12}$ | Cytoplasm/ER | [1] |
| $capsid_{empty}$ | $16\times10^{-12}$ | NPC | [1] |
| $capsid_{RNA}$ | $7.4\times10^{-12}$ | Cytoplasm/ER | [1] |
| $capsid_{RNA}$ | $16\times10^{-12}$ | NPC | [1] |
| $viral_{pol}$ | $22\times10^{-12}$ | Cytoplasm | Equals to L (assumption) |
| $RNA_{pol}$ | $22\times10^{-12}$ | Nucleus | Equals to $viral_{pol}$ (assumption) |
| $cccDNA.RNA_{pol}$ | $22\times10^{-12}$ | Nucleus | Equals to $viral_{pol}$ (assumption) |
| rcDNA | $0.07\times10^{-12}$ | NPC | [2] |
| rcDNA | $0.61\times10^{-12}$ | Nucleus | [2] |
| cccDNA | $0.61\times10^{-12}$ | Nucleus | [2] |
| LSmRNA | $0.61\times10^{-12}$ | Nucleus | [2] |
| pgRNA | $0.61\times10^{-12}$ | Nucleus | [2] |
| SmRNA | $0.61\times10^{-12}$ | Nucleus | [2] |

| | | | |
|---|---|---|---|
| XmRNA | $0.61\times10^{-12}$ | Nucleus | [2] |
| preCmRNA | $0.61\times10^{-12}$ | Nucleus | [2] |
| LSmRNA | $2.24\times10^{-12}$ | Cytoplasm | [2] |
| pgRNA | $2.24\times10^{-12}$ | Cytoplasm | [2] |
| SmRNA | $2.24\times10^{-12}$ | Cytoplasm | [2] |
| XmRNA | $2.24\times10^{-12}$ | Cytoplasm | [2] |
| preCmRNA | $2.24\times10^{-12}$ | Cytoplasm | [2] |
| LSmRNA | $0.07\times10^{-12}$ | NPC | [2] |
| pgRNA | $0.07\times10^{-12}$ | NPC | [2] |
| SmRNA | $0.07\times10^{-12}$ | NPC | [2] |
| XmRNA | $0.07\times10^{-12}$ | NPC | [2] |
| preCmRNA | $0.07\times10^{-12}$ | NPC | [2] |
| preC | $22\times10^{-12}$ | Cytoplasm | Equals to L (assumption) |
| S | $22\times10^{-12}$ | Cytoplasm/ER | Equals to L (assumption) |
| M | $22\times10^{-12}$ | Cytoplasm/ER | Equals to L (assumption) |
| L | $22\times10^{-12}$ | Cytoplasm/ER | scaled diffusion coefficient of GFP[3] based on [4] |
| X | $22\times10^{-12}$ | Cytoplasm | Equals to L (assumption) |
| interm | $9.7\times10^{-12}$ | Cytoplasm | ref [1] scaled based on [4] |
| RNP | $2.24\times10^{-12}$ | Cytoplasm | Equals to RNAs in cytoplasm |
| $capsid_{DNA}$.L(1,…20) | $22\times10^{-12}$ | ER | Equals to L (assumption) |
| $capsid_{RNA}$.L(1,…20) | $22\times10^{-12}$ | ER | Equals to L (assumption) |
| $capsid_{empty}$.L(1,…20) | $22\times10^{-12}$ | ER | Equals to L (assumption) |

Table S2. Reactions describing the degradation of different species that were included in the kinetic model

| Degradation reactions | Rate | Unit | Compartment |
|---|---|---|---|
| rcDNA → 0 | $1.60\times10^{-7}$ | $s^{-1}$ | Nucleus |
| cccDNA→ 0 | $1.60\times10^{-7}$ | $s^{-1}$ | Nucleus |
| pgRNA→ 0 | $3.8\times10^{-5}$ | $s^{-1}$ | Cytoplasm |
| LSmRNA→ 0 | $6.4\times10^{-5}$ | $s^{-1}$ | Cytoplasm |

| | | | |
|---|---|---|---|
| SmRNA→ 0 | $6.4\times10^{-5}$ | $s^{-1}$ | Cytoplasm |
| XmRNA→ 0 | $1.9\times10^{-4}$ | $s^{-1}$ | Cytoplasm |
| preCmRNA→ 0 | $3.8\times10^{-5}$ | $s^{-1}$ | Cytoplasm |
| L→ 0 | $2.9\times10^{-4}$ | $s^{-1}$ | Cytoplasm |
| M→ 0 | $2.9\times10^{-4}$ | $s^{-1}$ | Cytoplasm |
| S→ 0 | $2.9\times10^{-4}$ | $s^{-1}$ | Cytoplasm |
| dimer→ 0 | $2.9\times10^{-4}$ | $s^{-1}$ | Cytoplasm |
| $viral_{pol}$→ 0 | $2.9\times10^{-4}$ | $s^{-1}$ | Cytoplasm |
